# Tinnitus-like “hallucinations” elicited by sensory deprivation in an entropy maximization recurrent neural network

**DOI:** 10.1101/2021.01.11.426188

**Authors:** Aviv Dotan, Oren Shriki

## Abstract

Sensory deprivation has long been known to cause hallucinations or “phantom” sensations, the most common of which is tinnitus induced by hearing loss, affecting 10–20% of the population. An observable hearing loss, causing auditory sensory deprivation over a band of frequencies, is present in over 90% of people with tinnitus. Existing plasticity-based computational models for tinnitus are usually driven by homeostasis mechanisms, modeled to fit phenomenological findings. Here, we use an objective-driven learning algorithm to model an early auditory processing neuronal network, e.g., in the dorsal cochlear nucleus. The learning algorithm maximizes the network’s output entropy by learning the feed-forward and recurrent interactions in the model. We show that the connectivity patterns and responses learned by the model display several hallmarks of early auditory neuronal networks. We further demonstrate that attenuation of peripheral inputs drives the recurrent network towards its critical point and transition into a tinnitus-like state. In this state, the network activity resembles responses to genuine inputs even in the absence of external stimulation, namely, it “hallucinates” auditory responses. These findings demonstrate how objective-driven plasticity mechanisms that normally act to optimize the network’s input representation can also elicit pathologies such as tinnitus as a result of sensory deprivation.

**Author summary:** Tinnitus or “ringing in the ears” is a common pathology. It may result from mechanical damage in the inner ear, as well as from certain drugs such as salicylate (aspirin). A common approach toward a computational model for tinnitus is to use a neural network model with inherent plasticity applied to early auditory processing, where the input layer models the auditory nerve and the output layer models a nucleus in the brain stem. However, most of the existing computational models are phenomenological in nature, driven by a homeostatic principle. Here, we use an objective-driven learning algorithm based on information theory to learn the feed-forward interactions between the layers, as well as the recurrent interactions within the output layer. Through numerical simulations of the learning process, we show that attenuation of peripheral inputs drives the network into a tinnitus-like state, where the network activity resembles responses to genuine inputs even in the absence of external stimulation; namely, it “hallucinates” auditory responses. These findings demonstrate how plasticity mechanisms that normally act to optimize network performance can also lead to undesired outcomes, such as tinnitus, as a result of reduced peripheral hearing.

## Introduction

Tinnitus is a common form of auditory hallucinations, affecting the quality of life of many people (≈10–20% of the population, [1–6]). It can manifest as “ringing” in a narrow frequency band, but also as noise over a wide frequency range. An observable hearing loss, causing sensory deprivation over a band of frequencies, is present in *>*90% of people with tinnitus [1–4], and the remaining people with tinnitus are believed to suffer some damage in higher auditory processing pathways [5, 7] or have some cochlear damage that does not affect the audiogram [8].

From a neural processing point of view, hallucinations correspond to brain activity in sensory networks, which occurs in the absence of an objective external input. Hallucinations can occur in all sensory modalities, and can be induced by drugs, certain brain disorders, and sensory deprivation. For example, it is well known that visual deprivation (e.g., being in darkness for an extended period) elicits visual hallucinations, and, similarly, auditory deprivation elicits auditory hallucinations [9–11].

Although the causes of tinnitus can sometimes be mechanical (“objective tinnitus” [2, 12]), this is not the case in *>*95% of patients [6, 12]. This so-called “subjective tinnitus” is commonly associated with plasticity of feedback and recurrent neuronal circuits [2, 5, 8, 13–16].

The dorsal cochlear nucleus (DCN) is known to display tinnitus-related plastic reorganization following cochlear damage [17–20], and is thought to be a key player in the generation of tinnitus [21–24]. It is stimulated directly by the auditory nerve with a tonotopic mapping. Each output unit, composed of a group of different cells, receives inputs from a small number of input fibers and inhibits units of similar tuning [25, 26]. This connectivity pattern results in a sharp detection of specific notches [26]. As the DCN is the earliest candidate along the auditory path displaying tinnitus-related activity [17, 18], it is the most common candidate for the generation of tinnitus [21–24]. This choice is also supported by DCN hyperactivity following artificial induction of tinnitus [19, 20]. Interestingly, this induced hyperactivity persists even if the DCN is later isolated from inputs other than the auditory nerve [27]. This suggests that tinnitus-related hyperactivity in the DCN is intrinsic and not caused by feedback from higher order auditory networks.

While existing computational models successfully account for some of the characteristics of tinnitus [28], many of them are based on lateral inhibition [29–31] or gain adaptation [32], and do not take into account long-term neural plasticity. Plasticity-based models for tinnitus are usually phenomenological models, where plasticity is described as a homeostatic process [33–39] or an amplification of central noise [40], rather than as a process which serves a computational goal. Another computational model for tinnitus is based on stochastic resonance and suggests that tinnitus arises from an adaptive optimal noise level, but it is focused on a single auditory frequency and has yet to be further explored [41].

In this work, we try to gain new insights into tinnitus by using information theoretic-driven plasticity. We implemented the entropy maximization (EM) approach in a recurrent neural network [42] to model the connection between the raw sensory input and its downstream representation. This approach was previously applied to model the feed-forward connectivity in the primary visual cortex, giving rise to orientation-selective Gabor-like receptive fields [43]. A later generalization of the algorithm to learning recurrent connectivity [42] was used to show that EM drives recurrent visual neural networks toward critical behavior [44]. Furthermore, the evolved recurrent connectivity profile has a Mexican-hat shape; namely, neurons with similar preferred orientations tend to excite one another, while neurons with distant preferred orientations tend to inhibit one another. While the aforementioned studies focused on the normal function of the visual system, EM-based neural networks were barely used to model any abnormalities or to study the effect of changes in input statistics [45]. The relationship between EM-based adaptation and the emergence of tinnitus from sensory deprivation was previously discussed in the context of single neurons [46], yet it was never explored on a large-scale recurrent network.

Here, we trained a recurrent EM neural network to represent auditory stimuli, so it can stand as a simplified model for early auditory processing. Subsequently, to test the effect of sensory deprivation on the network’s output representation, we modified the input statistics by attenuating a certain frequency band. Our findings show that tinnitus-like hallucinations naturally arise in this model following sensory deprivation. These findings suggest that hallucinations following sensory deprivation can stem from general long-term plasticity mechanisms that act to optimize the representation of sensory information. Furthermore, our analysis indicates that the trained network tends to operate near a critical point on the verge of hallucinations, similar to previous findings [44]. The increased gain of the recurrent interactions, which acts to compensate for the attenuated input, may lead the network to cross the critical point into a regime of hallucinations.

## Results

To model the early stages of auditory processing (e.g., DCN), we used an entropy maximization (EM) approach to train a recurrent neural network (see Methods). The neurons obey first-order rate dynamics, and it is assumed that the network reaches a steady state following the presentation of each stimulus. The learning algorithm for the feed-forward and recurrent connectivity was based on the gradient-descent algorithm described in [42], with the addition of regularization. The network was trained in an unsupervised manner to represent simulated auditory stimuli (see Methods for more details). Figure 1 depicts the network’s architecture and a typical stimulus.

**Fig 1.**
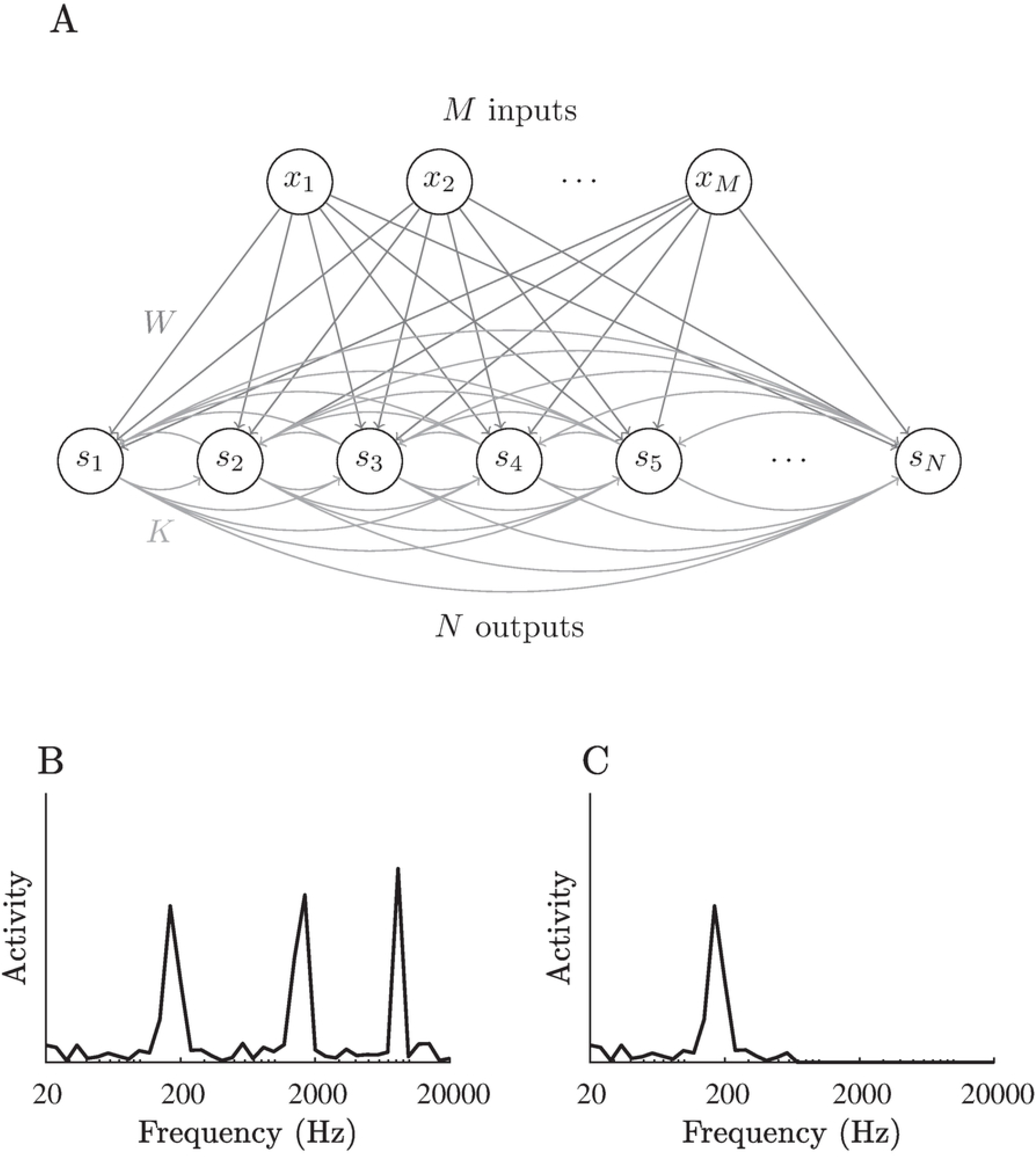
The neural network’s architecture and typical stimuli. A: The architecture of an overcomplete recurrent neural network. B: A typical simulated stimulus. C: The stimulus presented in B, after attenuation of high frequencies. Attenuation was achieved by multiplying the original input vector by an inverted sigmoidal function (see Methods).

In all simulations described here, we used a network of 40 input neurons and 400 output neurons (an overcomplete representation). Regularization was achieved using a cost on the norm of the weights and was applied to both feed-forward (using *ℓ*_1_ norm) and recurrent (using *ℓ*_2_ norm) sets of connections (see Methods). The coefficients of the regularization terms were set to *λ*_*W*_ = 0.001 for the feed-forward connections and *λ*_*K*_ = 0.183 for the recurrent connections (for details regarding these choices, see below the subsection on the Regularization effect).

### Training using typical stimuli

First, we trained the network using typical auditory inputs, simulated as a combination of multiple narrow Gaussians in the frequency domain with additional noise (see Methods and Fig. 1B). After the convergence of the learning process, each output neuron had a specific and unique preferred frequency (Fig. 2A). Furthermore, the connectivity profiles converged to a “Mexican-hat” shape for both feed-forward and recurrent connections (Fig. 2B,D). This profile of connectivity causes neurons with adjacent frequencies to excite one another, while neurons with slightly more distant frequencies inhibit each other. The significance of this profile lies in its ability to reduce the width of the output response profile for a Gaussian input, thus, effectively reducing the noise. Similarly shaped spectral receptive fields were observed in various primary auditory networks [25, 26, 47, 48] including the DCN, suggesting similar connectivity patterns.

**Fig 2.**
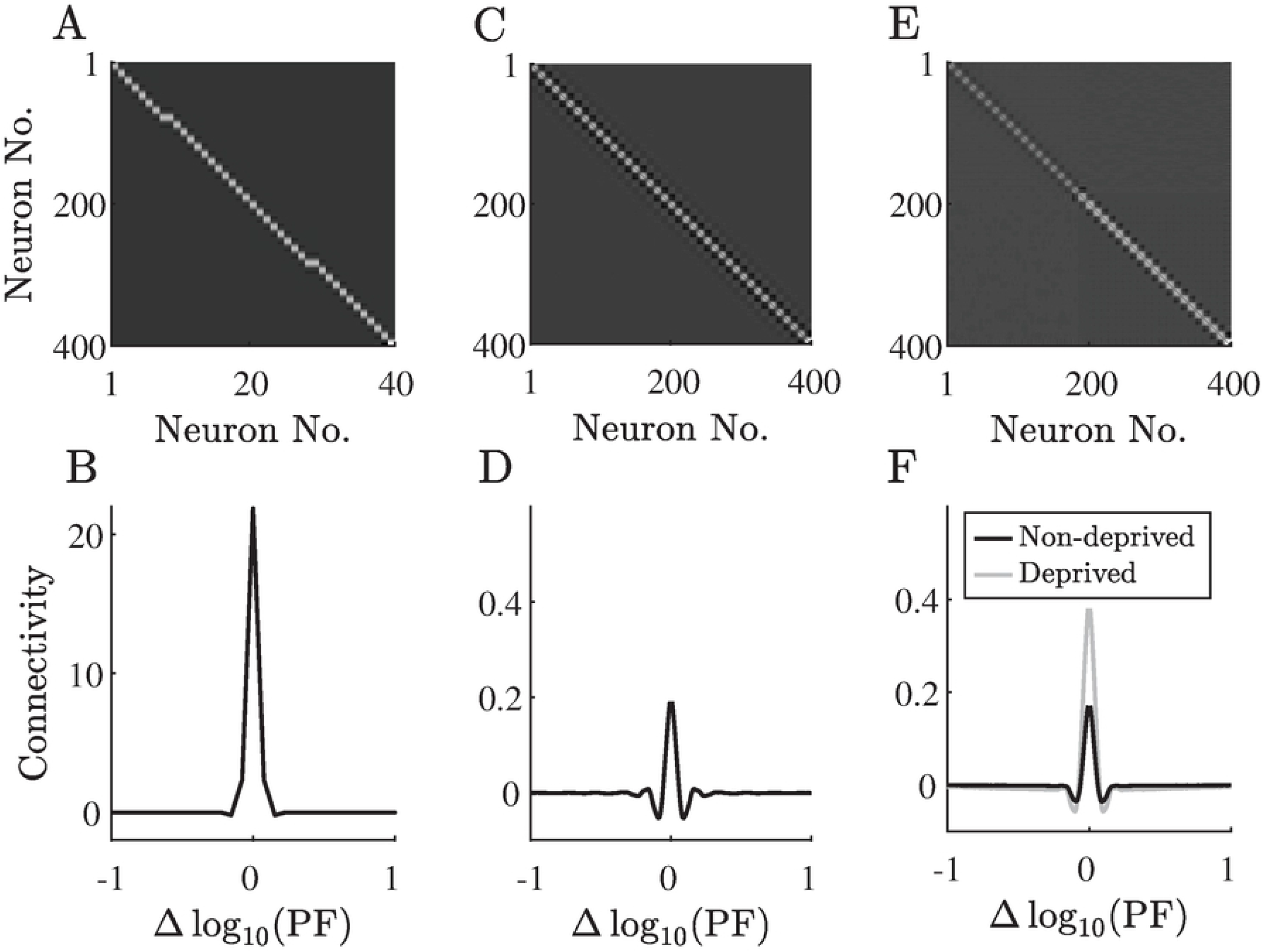
The network’s connectivity before and after sensory deprivation. A,B: The feed-forward connectivity matrix and its average row profile. C,D: The recurrent connectivity matrix and its average row profile before sensory deprivation. E,F: The recurrent connectivity matrix and its average row profiles after sensory deprivation, averaged separately for neurons in the deprived zone and the non-deprived zone. Each row profile is obtained by aligning the presynaptic connections to every neuron according to its preferred frequency and then averaging.

The network’s response to typical stimuli shows tonotopic responses, and the response in the absence of external stimuli is near spontaneous activity (Fig. 3A–F). We note that the initial feed-forward connectivity was manually tuned to produce a tonotopic mapping (using weak Gaussian profiles with ordered centers). Although the feed-forward connections do change throughout the learning process, the tonotopic organization remains stable. The tonotopic mapping is a well-known property of all auditory processing stages between the cochlea and the auditory cortex in various species, including humans [49–53]. The preservation of the tonotopic organization throughout the learning process is in agreement with biological observations, suggesting that it is created in the embryonic stages of development and is preserved through plasticity processes [54].

**Fig 3.**
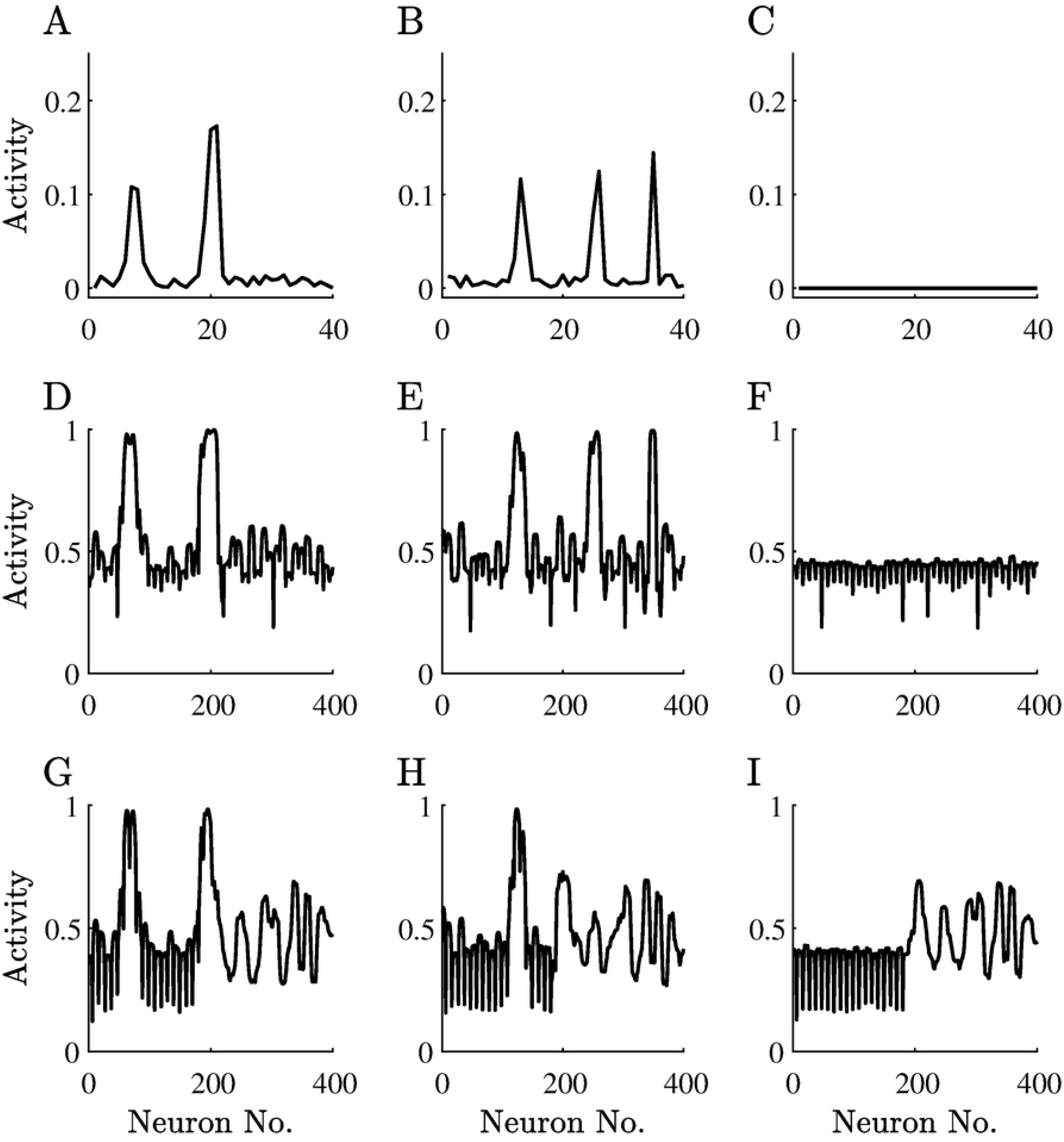
The network’s response to different stimuli before and after sensory deprivation. A,B: Typical stimuli. C: A silent stimulus (zero input). D–F: The network’s response to the stimuli presented in A–C. G–I: The network’s response to the stimuli presented in A–C after training on stimuli with attenuated high frequencies. Note that the spontaneous activity of the neurons is 0.5.

We noticed that spatial connectivity profiles barely change throughout the learning, while their scale changes dramatically. In light of this observation, we quantified several global parameters of the network as a function of the scale of the recurrent connectivity matrix (Fig. 4). We also used these measurements to gain insights into the effect of regularization on our results. First, note that the regularization caused the network learning process to converge to slightly down-scaled recurrent interactions compared to the optimal scale in terms of the non-regularized objective function (Fig. 4A). This specific scale seems to play a role in determining the proximity of the network dynamics to the critical point. Specifically, the convergence time rises dramatically at this point, reflecting the well-known phenomenon of “critical slowing down” [55–58]. In addition, at this scale, the population vector’s magnitude rises sharply, reflecting the emergence of non-uniform activity profiles in the absence of a structured input (see Methods and Fig. 4B,C). All these results point to the same conclusion – without the regularization, the recurrent connectivity should have been scaled by ≈ 1.3, such that the spectral radius of the recurrent connectivity matrix would be ≈4. We note that the maximal derivative of the chosen activation function 1*/*(1 + exp (−*x*)) is 1*/*4. Thus, having the spectral radius of the recurrent connectivity matrix near 4 indicates proximity to the critical point (see Methods). This means that the regularization keeps the recurrent connectivity below its optimal scale (in terms of the entropy term alone), and the network remains subcritical. We note that for different regularization coefficients, the scale of the interactions could obtain different values.

**Fig 4.**
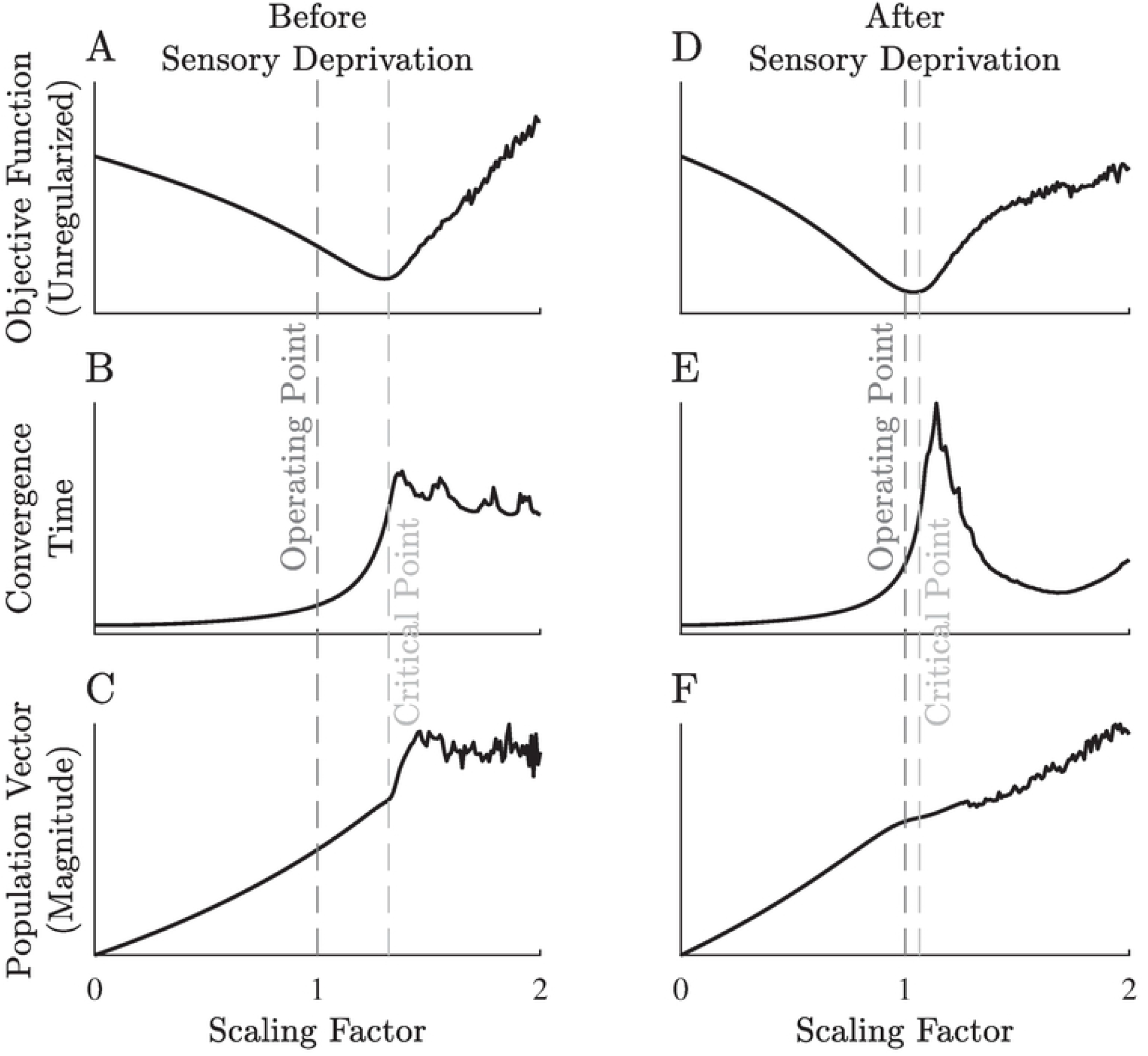
Global parameters for different scaling of the recurrent connections. A,D: The network’s objective function, without the regularization terms. B,E: The convergence time of the dynamics using Euler’s method. C,F: The population vector magnitude. All the above parameters are displayed for different scaling factors of the recurrent connectivity matrix, as found by the training process. In A–C, we used the recurrent connectivity matrix trained on typical stimuli, while in D–F, we used the recurrent connectivity matrix obtained after sensory deprivation. The operating point is at a scaling factor of 1, namely, the recurrent connectivity the learning process has converged to. The marked critical point (≈1.32 in A–C and ≈1.06 in D–F) is the scaling factor, for which the spectral radius of the recurrent connectivity matrix is 4. For visualization purposes, the population vector magnitudes (C,F) are displayed on different scales.

### Sensory deprivation

After the learning was stabilized for normal stimuli, we attenuated the inputs in the higher half of the frequency range (Fig. 1C), and let the network’s recurrent connections adapt to the new input statistics. Consequently, the recurrent connectivity between the deprived neurons was strengthened (Fig. 2E,F). The stronger recurrent connectivity in the deprived region led to a phase transition, resulting in an inhomogeneous stationary activity pattern independent of the given input (Fig. 3G–I). We interpret those results as “hallucinations”, elicited by the sensory deprivation. Interestingly, the “hallucinations” in our model develop only in the deprived region of the output layer, consistent with certain types of tinnitus [3, 46, 59, 60].

Following the induction of sensory deprivation, we evaluated the criticality measures once again (Fig. 4D–F). The results remained qualitatively similar, but the optimal scale moved much closer to 1 (≈1.06). Thus, the network converged to a point much closer to its critical point, compared to its state before the induction of sensory deprivation. We note that following sensory deprivation, the effect of learning on the recurrent connections is not limited to scaling. Hence, the different measures exhibit different patterns in the supercritical domain (above the scale of ≈1.06).

### Regularization effect

As discussed above, to keep the dynamics from crossing into the supercritical domain, we added regularization to the network’s weights. For each type of connectivity matrix (feed-forward and recurrent), we tested regularization both by *ℓ*_1_ and *ℓ*_2_ norms of the connections. Applying *ℓ*_1_ regularization is known to lead to sparse connectivity [61]; however, applying it to the recurrent connectivity matrix ended in nullifying all connections but two, which were still strong enough to turn the dynamics into the supercritical domain. These results are extremely non-biological (as recurrent connectivity is present in most biological neural networks); thus, we focus only on simulations where the recurrent connections were regularized by their *ℓ*_2_ norm. Using either the *ℓ*_1_ or *ℓ*_2_ norm to regularize the feed-forward connectivity did not have a dramatic effect on the results. Since using the *ℓ*_1_ norm leads to a more biological sparse connectivity, as found experimentally in the DCN [26], we chose to focus on this option.

The stability of the network’s fixed point is determined by the sign of the eigenvalues of the matrix that controls the linearized dynamics. In this case, the corresponding matrix is (*I* − *GK*), where *K* is the recurrent connectivity matrix and *G* is a diagonal matrix containing the derivatives of the activation function for each output neuron (see Methods). Since the maximal derivative of the chosen activation function (1*/*(1 + exp (−*x*))) is 1*/*4, the critical point is characterized by having the spectral radius of the recurrent connectivity matrix, *K*, near 4. We used this result as an efficient surrogate to the actual critical point.

In our simulations, the spectral radius of the recurrent connectivity matrix *K* decreased with the respective regularization coefficient *λ*_*K*_, with a characteristic sharp drop (Fig. 5). Generally, the value of *λ*_*K*_ where this drop occurs depends mainly on the number of output neurons; however, in our simulations, sensory deprivation caused this value to rise. This phenomenon created an interval of *λ*_*K*_ values, where sensory deprivation drives the dynamics much closer to the critical point, thus, eliciting the hallucination-like responses described before. To emphasize this effect, we used the lower bound of this interval (*λ*_*K*_=0.183) in all simulations previously displayed; for larger values of *λ*_*K*_, the system will be further away from the critical point.

**Fig 5.**
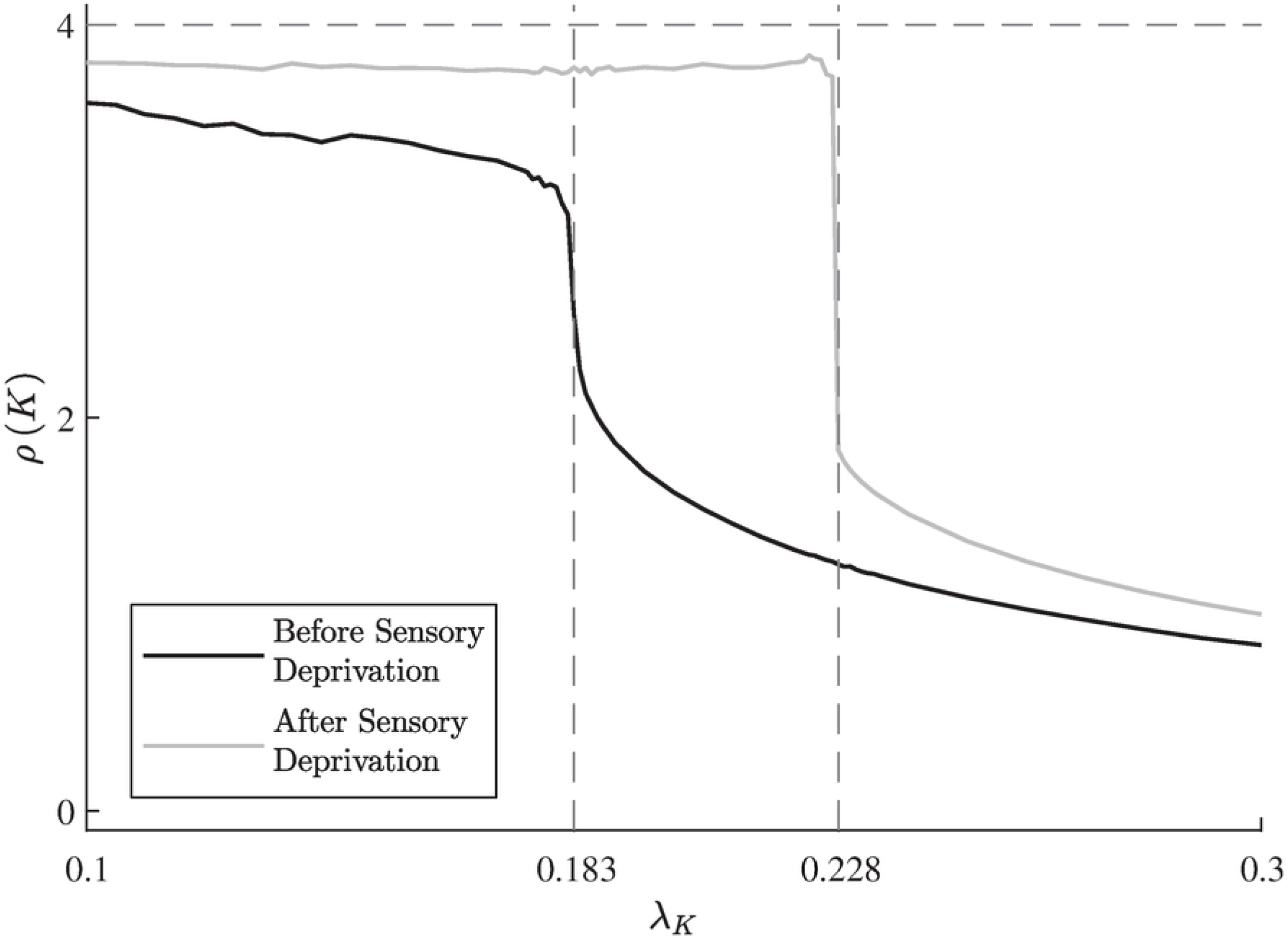
Regularization effect on the spectral radius of the recurrent connectivity matrix. The spectral radius of the recurrent connectivity matrix *K* decreases with the regularization coefficient *λ*_*K*_, before and after the induction of sensory deprivation. Due to the chosen sigmoidal activation function, the sharp drop in the spectral radius from ≈4 to ≈2 determines the border between near-critical and subcritical dynamics. After the induction of sensory deprivation, this border moves to higher values of the regularization coefficient, hence, creating an interval (from ≈0.183 to ≈0.228) of regularization coefficient values where sensory deprivation causes “hallucinations”.

## Discussion

In this work, we used an EM approach to train a recurrent neural network to represent simulated auditory stimuli, and examined the effect of input statistics on the evolved representation. For typical inputs, the network developed connectivity patterns and exhibited output responses similar to biological findings regarding the auditory system in general [62–65] and, more specifically, the DCN [25, 26]. Interestingly, sensory deprivation elicited tinnitus-like “hallucinations” in the network, resembling the characteristics of certain types of tinnitus [3, 11, 46, 59, 60]. Although we focused here on tinnitus, this qualitative phenomenon is independent of the input modality and can be used to explain how other kinds of “phantom” sensations are caused by neural plasticity and involve the specific region in the sensory input space, which was deprived of input [66, 67].

Previous computational models relied on phenomenological homeostasis-driven plasticity to demonstrate tinnitus elicited by sensory deprivation [33–38]. Here, we used an objective-driven plasticity, namely, the main mechanism underlying the network’s plasticity is optimizing an explicit computational goal. Specifically, the network maximizes the entropy of its output, which corresponds to increasing input sensitivity [44]. The general resemblance of our model to biological findings supports the hypothesis that EM serves as a computational objective for primary sensory processing networks in the brain (e.g., [43, 44]). However, as described in the Methods section, the vanilla EM learning rules drive the network into a phase transition. This process leads the network away from a stable fixed point and into dynamical states with poor information representation. Thus, some regularization should be used to keep the network subcritical. To this end, we used a penalty on the *ℓ*_2_ norm of the recurrent connections as a regularization method, which can be thought of as a kind of homeostatic mechanism [68–72]. In this model, the emergence of tinnitus depends on the interplay between the computational objective and the homeostatic regularization, in contrast to models driven by a single phenomenological homeostatic mechanism. Future studies might employ different types of regularization methods (e.g., firing-rate-based rather than weight-based) and examine their effect on the dynamics of the network.

While most of the hyper-parameters of the model can be chosen arbitrarily without having any qualitative effect on the results, the regularization coefficient for the recurrent connectivity, *λ*_*K*_, is an exception; if it is too small, numerical instabilities might accidentally drive the network into a supercritical domain, but if it is too large, the network will always remain subcritical. In the first case, the output may no longer be dependent on the input, while in the second case, the the input may have little effect on the output – in both cases, moving away from the critical point leads to poor sensitivity. In practice, there is a specific range of values which yields the qualitative results demonstrated in this paper (see Fig. 5) and, according to our observations, it is mainly dependent on the number of output neurons. Here, we used a grid search to find the corresponding range, and the results were obtained using the minimal value within it. This choice minimized the cost of regularization relative to the EM objective, while keeping the evolved dynamics in the subcritical regime. Furthermore, this choice of *λ*_*K*_ drives the network close to the critical point and emphasizes the effect of sensory deprivation on the transition into the tinnitus-like domain and on the resulting “hallucinations”.

These results are in line with a plethora of studies from recent years, suggesting near-critical dynamics in biological neural networks across various scales, from neuronal cultures to large-scale human brain activity [73–81]. In particular, it is proposed that healthy neural dynamics are poised near a critical point, yet within the subcritical domain [82]. Under these circumstances, changes in the input statistics can trigger the network to transition into supercritical dynamics, which may manifest as hallucinations. Our study portrays a concrete, albeit simplified, network model that leads to near-critical dynamics and experiences a transition from healthy to pathological neural dynamics as a consequence of inherent plasticity and sensory deprivation. We note that the network dynamics here are too simplified to enable a direct comparison with the rich dynamics observed in cortical networks and with common hallmarks of criticality (e.g., [73]).

An illuminating perspective on the emergence of hallucinations, such as tinnitus, as a consequence of sensory deprivation comes from the framework of Bayesian inference [83–85]. According to this framework, sensory systems generate perception by combining the incoming stimuli with prior expectations in a way that takes into account the relative uncertainty of each. Under sensory deprivation, the uncertainty about the input is very large; hence, the weight of the prior expectations become more dominant. This process may eventually lead to a state in which prior expectations dominate perception, which can be interpreted as a hallucination [86]. If this perception is maintained long enough, it will turn into a strong prior by itself, thus, giving rise to a chronic hallucination – namely, tinnitus [84]. Although our model does not use the Bayesian framework explicitly, it does share a few characteristics with the Bayesian approach. For example, according to the Bayesian framework, the profile of the “hallucination” is an amplified prior, so it should resemble typical inputs–much like our results (see Fig. 3G–I). The advantage of the model described here lies in its mechanistic nature, namely, that it is cast in the language of neuronal networks with long-term plasticity. Thus, it can be more straightforward to interpret and compare to experimental data.

To summarize, we have demonstrated how the EM approach can be used as a model for early auditory processing and the phenomenon of tinnitus. Previous works have demonstrated that EM-based neural networks can serve as models for early visual processing [43, 44] and the phenomenon of synaesthesia [45]. We believe that this framework can be used for modeling other modalities and phenomena as well. It is also important to extend this framework to more biologically plausible network models, which could account for more detailed aspects of the underlying neural dynamics.

## Methods

### The model

We modeled an early auditory processing neural network (e.g., the DCN) using the overcomplete recurrent EM neural network described in [42], with the addition of regularization on strong connectivity.

### Network architecture and dynamics

Our system is composed of *M* input neurons, **x**, and *N* output neurons, **s**. Each output neuron’s activity through time is given by the dynamic equation:

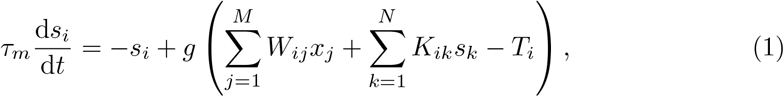

where *W* is the feed-forward connectivity matrix, *K* is the recurrent connectivity matrix, *T* are the output neurons’ thresholds, and *g* (*x*) = 1*/*(1 + exp (−*x*)) is the activation function of the neurons. For overcomplete transformations, we assume *M < N* (Fig. 1A).

The fixed points of Eq. 1 are given implicitly by:

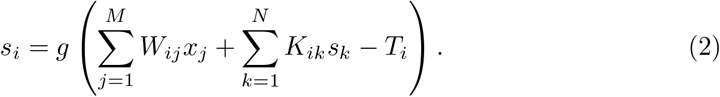

These fixed points are stable iff all of the eigenvalues of the linearized dynamics matrix (*I* − *GK*) have positive real parts [44] [*G* is a diagonal matrix defined by 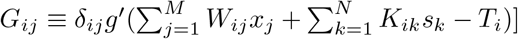. Since the values of *G* are upper-bounded by max_*x*_ *g* (*x*) = 1*/*4, for a matrix *K* with eigenvalues *<*4, the fixed points are indeed stable. In practice, when fixed points exist at all, there will usually only be one such stable fixed point.

Numerically, the steady state can be found via integrating Eq. 1 using Euler’s method for a long time-period until the activities stabilize; however, this method is highly inefficient. In this work, we found the steady state by solving Eq. 2 directly using the Newton-Raphson method.

When the eigenvalues of *K* are near 4, the eigenvalues of (*I* − *GK*) might get close to zero. Crossing this point will result in instability of the fixed point and a phase transition. Near this phase transition, the decrease in the eigenvalues of (*I* − *GK*) will cause the effective time constant to rise – a phenomenon termed “critical slowing down”. Furthermore, such a phase transition is expected to be characterized by a spontaneous symmetry breaking [87], which can be measured by several metrics. Here, we used the population vector for that purpose, calculated as 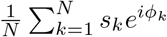 where *ϕ*_*k*_ ≡ 2*πk/N* . Although in our case the boundary conditions are not periodic, we assume their effect to be negligible since *N ≫* 1.

### Learning rules

The goal of the network is to find the set {*W* ^∗^, *K*^∗^, *T* ^∗^}which maximizes the entropy *H*(**s**) of the steady state outputs. To do so, we used the objective function described in [42], with additional regularization terms on the *ℓ*_1_ and *ℓ*_2_ norms of *W* and *K*, respectively:

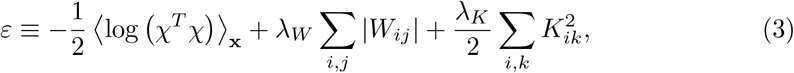

where 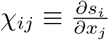 is the Jacobian of the transformation given by *χ* = *ϕW* , and *ϕ* ≡ − (*I* − *GK*)^−1^*G* [42].

This objective function, without the regularization terms, would lead to an increase in the singular values of *χ*. One way to achieve that goal is to decrease the eigenvalues of (*I GK*) to zero, which may lead one of them to turn slightly negative due to numerical errors. This will result in instability of the fixed point and a phase transition, as discussed above. The goal of the regularization terms is to prevent this phenomenon, which is a general property of unregularized entropy maximization systems of continuous variables [88].

The learning rules were derived using the gradient descent method, as in [42]:

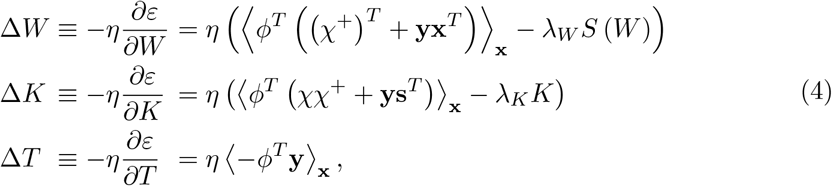

where 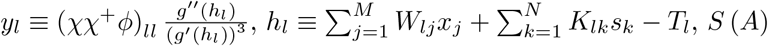 is defined by (*S* (*A*)) sign (*A*_*ij*_) and *χ*^+^ stands for the pseudo-inverse of *χ* (in the overcomplete case used here, *χ*^+^ = *χ*^*T*^ *χ χ*^*T*^).

### Auditory inputs

The input stimuli were chosen according to certain heuristics to emulate the system’s response to tones of varying frequencies and amplitudes. Each input sample embodies the reaction of the auditory hair cells to a combination of tones of certain frequencies. The input profile for a pure tone is centered on the neuron that best matches that frequency, and drops off to neighboring neurons to form a narrow Gaussian response curve. The amplitude and the frequency were selected at random with a uniform distribution on the permitted ranges. In addition to the input response, all neurons feature some spontaneous activity that is irrespective of the inputs to model the neurons’ reaction to background noises and non-stimulated motion of the hair cells (Fig. 1B).

The amplitudes of natural sounds are not uniformly distributed, loud sounds being exponentially less common; however, the response of the inner hair cells is determined not only by the absolute amplitude of the sound, but also by the reactivity of the basilar membrane, as controlled by the outer hair cells. This serves as an automatic gain control mechanism, giving the inner hair cells use of their full motion capacity for normal inputs. Therefore, we hold the uniform distribution to be a good approximation to the output of the inner hair cells when presented with natural sounds [89, 90] To model sensory deprivation, we attenuated the higher half of the frequency domain by applying a (monotonically decreasing) sigmoid envelope to all stimuli (Fig. 1C). The choice of attenuating the higher frequencies was based on the most common type of hearing loss [91, 92], but attenuation could be applied to other frequency bands.

## Acknowledgments

The authors wish to thank Avishalom Shalit and Jennifer Resnik for helpful discussions and valuable comments on the manuscript.

## References

1. Nicolas-Puel C, Akbaraly T, Lloyd R, Berr C, Uziel A, Rebillard G, et al. Characteristics of tinnitus in a population of 555 patients: Specificities of tinnitus induced by noise trauma. The International Tinnitus Journal. 2006;12(1):64–70.

2. Saunders JC. The role of central nervous system plasticity in tinnitus. Journal of Communication Disorders. 2007;40(4):313–334. doi:10.1016/j.jcomdis.2007.03.006.

3. Roberts LE, Eggermont JJ, Caspary DM, Shore SE, Melcher JR, Kaltenbach JA. Ringing ears: The neuroscience of tinnitus. Journal of Neuroscience. 2010;30(45):14972–14979. doi:10.1523/jneurosci.4028-10.2010.

4. Shargorodsky J, Curhan GC, Farwell WR. Prevalence and characteristics of tinnitus among US adults. The American Journal of Medicine. 2010;123(8):711–718. doi:10.1016/j.amjmed.2010.02.015.

5. Langguth B, Kreuzer PM, Kleinjung T, Ridder DD. Tinnitus: Causes and clinical management. The Lancet Neurology. 2013;12(9):920–930. doi:10.1016/S1474-4422(13)70160-1.

6. McCormack A, Edmondson-Jones M, Somerset S, Hall D. A systematic review of the reporting of tinnitus prevalence and severity. Hearing Research. 2016;337:70–79. doi:10.1016/j.heares.2016.05.009.

7. Weisz N, Hartmann T, Dohrmann K, Schlee W, Noreña A. High-frequency tinnitus without hearing loss does not mean absence of deafferentation. Hearing Research. 2006;222(1):108–114. doi:10.1016/j.heares.2006.09.003.

8. Noreña AJ, Farley BJ. Tinnitus-related neural activity: Theories of generation, propagation, and centralization. Hearing Research. 2013;295:161–171. doi:10.1016/j.heares.2012.09.010.

9. Zuckerman M, Cohen N. Sources of reports of visual and auditory sensations in perceptual-isolation experiments. Psychological Bulletin. 1964;62(1):1–20. doi:10.1037/h0048599.

10. Merabet LB, Maguire D, Warde A, Alterescu K, Stickgold R, Pascual-Leone A. Visual hallucinations during prolonged blindfolding in sighted subjects. Journal of Neuro-Ophthalmology. 2004;24(2). doi:10.1097/00041327-200406000-00003.

11. Schaette R, Turtle C, Munro KJ. Reversible induction of phantom auditory sensations through simulated unilateral hearing loss. PLoS One. 2012;7(6):1–6. doi:10.1371/journal.pone.0035238.

12. Blom JD, Sommer IEC. Auditory hallucinations: nomenclature and classification. Cognitive and Behavioral Neurology. 2010;23(1).

13. Eggermont JJ, Roberts LE. The neuroscience of tinnitus. Trends in Neurosciences. 2004;27(11):676–682. doi:10.1093/acprof:oso/9780199605606.001.0001.

14. Rauschecker JP, Leaver AM, Mühlau M. Tuning out the noise: Limbic-auditory interactions in tinnitus. Neuron. 2010;66(6):819–826. doi:10.1016/j.neuron.2010.04.032.

15. Wang H, Brozoski TJ, Caspary DM. Inhibitory neurotransmission in animal models of tinnitus: Maladaptive plasticity. Hearing Research. 2011;279(1):111–117. doi:10.1016/j.heares.2011.04.004.

16. Henry JA, Roberts LE, Caspary DM, Theodoroff SM, Salvi RJ. Underlying mechanisms of tinnitus: Review and clinical implications. Journal of the American Academy of Audiology. 2014;25(01):5–22. doi:10.3766/jaaa.25.1.2.

17. Brozoski TJ, Bauer CA, Caspary DM. Elevated fusiform cell activity in the dorsal cochlear nucleus of chinchillas with psychophysical evidence of tinnitus. Journal of Neuroscience. 2002;22(6):2383–2390. doi:10.1523/jneurosci.22-06-02383.2002.

18. Brozoski TJ, Bauer CA. The effect of dorsal cochlear nucleus ablation on tinnitus in rats. Hearing Research. 2005;206(1):227–236. doi:10.1016/j.heares.2004.12.013.

19. Wang H, Brozoski TJ, Turner JG, Ling L, Parrish JL, Hughes LF, et al. Plasticity at glycinergic synapses in dorsal cochlear nucleus of rats with behavioral evidence of tinnitus. Neuroscience. 2009;164(2):747–759. doi:10.1016/j.neuroscience.2009.08.026.

20. Dehmel S, Pradhan S, Koehler S, Bledsoe S, Shore S. Noise overexposure alters long-term somatosensory-auditory processing in the dorsal cochlear nucleus—Possible basis for tinnitus-related hyperactivity? Journal of Neuroscience. 2012;32(5):1660–1671. doi:10.1523/jneurosci.4608-11.2012.

21. Kaltenbach JA. The dorsal cochlear nucleus as a participant in the auditory, attentional and emotional components of tinnitus. Hearing Research. 2006;216–217:224–234. doi:10.1016/j.heares.2006.01.002.

22. Kaltenbach JA, Godfrey DA. Dorsal cochlear nucleus hyperactivity and tinnitus: Are they related? American Journal of Audiology. 2008;17(2):S148–S161. doi:10.1044/1059-0889(2008/08-0004).

23. Tzounopoulos T. Mechanisms of synaptic plasticity in the dorsal cochlear nucleus: Plasticity-induced changes that could underlie tinnitus. American Journal of Audiology. 2008;17(2):S170–S175. doi:10.1044/1059-0889(2008/07-0030).

24. Baizer JS, Manohar S, Paolone NA, Weinstock N, Salvi RJ. Understanding tinnitus: The dorsal cochlear nucleus, organization and plasticity. Brain Research. 2012;1485:40–53. doi:10.1016/j.brainres.2012.03.044.

25. Spirou GA, Davis KA, Nelken I, Young ED. Spectral integration by Type II interneurons in dorsal cochlear nucleus. Journal of Neurophysiology. 1999;82(2):648–663. doi:10.1152/jn.1999.82.2.648.

26. Oertel D, Young ED. What’s a cerebellar circuit doing in the auditory system? Trends in Neurosciences. 2004;27(2):104–110. doi:10.1016/j.tins.2003.12.001.

27. Zhang JS, Kaltenbach JA, Godfrey DA, Wang J. Origin of hyperactivity in the hamster dorsal cochlear nucleus following intense sound exposure. Journal of Neuroscience Research. 2006;84(4):819–831. doi:10.1002/jnr.20985.

28. Schaette R, Kempter R. Computational models of neurophysiological correlates of tinnitus. Frontiers in Systems Neuroscience. 2012;6:34. doi:10.3389/fnsys.2012.00034.

29. Gerken GM. Central tinnitus and lateral inhibition: An auditory brainstem model. Hearing Research. 1996;97(1):75–83. doi:10.1016/S0378-5955(96)80009-8.

30. Kral A, Majernik V. On lateral inhibition in the auditory system. General Physiology and Biophysics. 1996;15(2):109–127.

31. Bruce IC, Bajaj HS, Ko J. Lateral-inhibitory-network models of tinnitus. IFAC Proceedings Volumes. 2003;36(15):359–363. doi:10.1016/S1474-6670(17)33529-2.

32. Parra LC, Pearlmutter BA. Illusory percepts from auditory adaptation. The Journal of the Acoustical Society of America. 2007;121(3):1632–1641. doi:10.1121/1.2431346.

33. Dominguez M, Becker S, Bruce I, Read H. A spiking neuron model of cortical correlates of sensorineural hearing loss: Spontaneous firing, synchrony, and tinnitus. Neural Computation. 2006;18(12):2942–2958. doi:10.1162/neco.2006.18.12.2942.

34. Schaette R, Kempter R. Development of tinnitus-related neuronal hyperactivity through homeostatic plasticity after hearing loss: a computational model. European Journal of Neuroscience. 2006;23(11):3124–3138. doi:10.1111/j.1460-9568.2006.04774.x.

35. Schaette R, Kempter R. Predicting tinnitus pitch from patients’ audiograms with a computational model for the development of neuronal hyperactivity. Journal of Neurophysiology. 2009;101(6):3042–3052. doi:10.1152/jn.91256.2008.

36. Schaette R, McAlpine D. Tinnitus with a normal audiogram: Physiological evidence for hidden hearing loss and computational model. Journal of Neuroscience. 2011;31(38):13452–13457. doi:10.1523/jneurosci.2156-11.2011.

37. Chrostowski M, Yang L, Wilson HR, Bruce IC, Becker S. Can homeostatic plasticity in deafferented primary auditory cortex lead to travelling waves of excitation? Journal of Computational Neuroscience. 2011;30(2):279–299. doi:10.1007/s10827-010-0256-1.

38. Gault R, McGinnity TM, Coleman S. A computational model of thalamocortical dysrhythmia in tinnitus sufferers. IEEE Transactions on Neural Systems and Rehabilitation Engineering. 2018;doi:10.1109/tnsre.2018.2863740.

39. Gault R, McGinnity TM, Coleman S. Perceptual modeling of tinnitus pitch and loudness. IEEE Transactions on Cognitive and Developmental Systems. 2020;12(2):332–343. doi:10.1109/tcds.2020.2964841.

40. Zeng FG. An active loudness model suggesting tinnitus as increased central noise and hyperacusis as increased nonlinear gain. Hearing Research. 2013;295:172–179. doi:10.1016/j.heares.2012.05.009.

41. Krauss P, Tziridis K, Metzner C, Schilling A, Hoppe U, Schulze H. Stochastic resonance controlled upregulation of internal noise after hearing loss as a putative cause of tinnitus-related neuronal hyperactivity. Frontiers in Neuroscience. 2016;10:597. doi:10.3389/fnins.2016.00597.

42. Shriki O, Sompolinsky H, Lee DD. An information maximization approach to overcomplete and recurrent representations. In: Leen TK, Dietterich TG, Tresp V, editors. Advances in Neural Information Processing Systems 13. MIT Press; 2001. p. 612–618.

43. Bell AJ, Sejnowski TJ. The “independent components” of natural scenes are edge filters. Vision Research. 1997;37(23):3327–3338. doi:10.1016/S0042-6989(97)00121-1.

44. Shriki O, Yellin D. Optimal information representation and criticality in an adaptive sensory recurrent neuronal network. PLoS Comput Biol. 2016;12(2):e1004698. doi:10.1371/journal.pcbi.1004698.

45. Shriki O, Sadeh Y, Ward J. The emergence of synaesthesia in a neuronal network model via changes in perceptual sensitivity and plasticity. PLoS Comput Biol. 2016;12(7):e1004959. doi:10.1371/journal.pcbi.1004959.

46. Noreña AJ. An integrative model of tinnitus based on a central gain controlling neural sensitivity. Neuroscience & Biobehavioral Reviews. 2011;35(5):1089–1109. doi:10.1016/j.neubiorev.2010.11.003.

47. Depireux DA, Simon JZ, Klein DJ, Shamma SA. Spectro-temporal response field characterization with dynamic ripples in ferret primary auditory cortex. Journal of Neurophysiology. 2001;85(3):1220–1234. doi:10.1152/jn.2001.85.3.1220.

48. Nagel KI, Doupe AJ. Organizing principles of spectro-temporal encoding in the avian primary auditory area field L. Neuron. 2008;58(6):938–955. doi:10.1016/j.neuron.2008.04.028.

49. Rubel E, Parks T. Organization and development of brain stem auditory nuclei of the chicken: Tonotopic organization of n. magnocellularis and n. laminaris. The Journal of Comparative Neurology. 1975;164(4):411–433. doi:10.1002/cne.901640403.

50. Yu X, Sanes DH, Aristizabal O, Wadghiri YZ, Turnbull DH. Large-scale reorganization of the tonotopic map in mouse auditory midbrain revealed by MRI. Proceedings of the National Academy of Sciences. 2007;104(29):12193–12198. doi:10.1073/pnas.0700960104.

51. Morel A, Garraghty PE, Kaas JH. Tonotopic organization, architectonic fields, and connections of auditory cortex in macaque monkeys. J Comp Neurol. 1993;335(3):437–459. doi:10.1002/cne.903350312.

52. Romani GL, Williamson SJ, Kaufman L. Tonotopic organization of the human auditory cortex. Science. 1982;216(4552):1339–1340. doi:10.1126/science.7079770.

53. Formisano E, Kim DS, Di Salle F, van de Moortele PF, Ugurbil K, Goebel R. Mirror-symmetric tonotopic maps in human primary auditory cortex. Neuron. 2003;40(4):859–869. doi:10.1016/S0896-6273(03)00669-X.

54. Kandler K, Clause A, Noh J. Tonotopic reorganization of developing auditory brainstem circuits. Nature Neuroscience. 2009;12(6):711–717. doi:10.1038/nn.2332.

55. Hohenberg PC, Halperin BI. Theory of dynamic critical phenomena. Rev Mod Phys. 1977;49:435–479. doi:10.1103/RevModPhys.49.435.

56. Strogatz SH. Nonlinear dynamics and chaos: with applications to physics, biology, chemistry, and engineering. CRC Press; 2018.

57. Djurberg C, Svedlindh P, Nordblad P, Hansen MF, Bødker F, Mørup S. Dynamics of an interacting particle system: Evidence of critical slowing down. Phys Rev Lett. 1997;79:5154–5157. doi:10.1103/PhysRevLett.79.5154.

58. Dai L, Vorselen D, Korolev KS, Gore J. Generic indicators for loss of resilience before a tipping point leading to population collapse. Science. 2012;336(6085):1175–1177. doi:10.1126/science.1219805.

59. Noreña A, Micheyl C, Chery-Croze S, Collet L. Psychoacoustic characterization of the tinnitus spectrum: Implications for the underlying mechanisms of tinnitus. Audiology and Neurotology. 2002;7(6):358–369. doi:10.1159/000066156.

60. König O, Schaette R, Kempter R, Gross M. Course of hearing loss and occurrence of tinnitus. Hearing Research. 2006;221(1):59–64. doi:10.1016/j.heares.2006.07.007.

61. Hastie T, Tibshirani R, Friedman J. The elements of statistical learning: data mining, inference, and prediction. 2nd ed. Springer Series in Statistics. Springer, New York, NY; 2009.

62. Reale RA, Imig TJ. Tonotopic organization in auditory cortex of the cat. Journal of Comparative Neurology. 1980;192(2):265–291. doi:10.1002/cne.901920207.

63. Escabí MA, Schreiner CE. Nonlinear spectrotemporal sound analysis by neurons in the auditory midbrain. Journal of Neuroscience. 2002;22(10):4114–4131. doi:10.1523/jneurosci.22-10-04114.2002.

64. Humphries C, Liebenthal E, Binder JR. Tonotopic organization of human auditory cortex. NeuroImage. 2010;50(3):1202–1211. doi:10.1016/j.neuroimage.2010.01.046.

65. Levy RB, Reyes AD. Spatial profile of excitatory and inhibitory synaptic connectivity in mouse primary auditory cortex. Journal of Neuroscience. 2012;32(16):5609–5619. doi:10.1523/jneurosci.5158-11.2012.

66. Pons TP, Garraghty PE, Ommaya AK, Kaas JH, Taub E, Mishkin M. Massive cortical reorganization after sensory deafferentation in adult macaques. Science. 1991;252(5014):1857–1861. doi:10.1126/science.1843843.

67. Grüsser SM, Mühlnickel W, Schaefer M, Villringer K, Christmann C, Koeppe C, et al. Remote activation of referred phantom sensation and cortical reorganization in human upper extremity amputees. Experimental Brain Research. 2004;154(1):97–102. doi:10.1007/s00221-003-1649-4.

68. Turrigiano GG, Leslie KR, Desai NS, Rutherford LC, Nelson SB. Activity-dependent scaling of quantal amplitude in neocortical neurons. Nature. 1998;391(6670):892–896. doi:10.1038/36103.

69. Turrigiano GG, Nelson SB. Hebb and homeostasis in neuronal plasticity. Current Opinion in Neurobiology. 2000;10(3):358–364. doi:10.1016/S0959-4388(00)00091-X.

70. Turrigiano GG, Nelson SB. Homeostatic plasticity in the developing nervous system. Nature Reviews Neuroscience. 2004;5(2):97–107. doi:10.1038/nrn1327.

71. Turrigiano GG. The self-tuning neuron: Synaptic scaling of excitatory synapses. Cell. 2008;135(3):422–435. doi:10.1016/j.cell.2008.10.008.

72. Styr B, Slutsky I. Imbalance between firing homeostasis and synaptic plasticity drives early-phase Alzheimer’s disease. Nature Neuroscience. 2018;21(4):463–473. doi:10.1038/s41593-018-0080-x.

73. Beggs JM, Plenz D. Neuronal avalanches in neocortical circuits. Journal of Neuroscience. 2003;23(35):11167–11177. doi:10.1523/jneurosci.23-35-11167.2003.

74. Tagliazucchi E, Balenzuela P, Fraiman D, Chialvo D. Criticality in large-scale brain fMRI dynamics unveiled by a novel point process analysis. Frontiers in Physiology. 2012;3:15. doi:10.3389/fphys.2012.00015.

75. Palva JM, Zhigalov A, Hirvonen J, Korhonen O, Linkenkaer-Hansen K, Palva S. Neuronal long-range temporal correlations and avalanche dynamics are correlated with behavioral scaling laws. Proceedings of the National Academy of Sciences. 2013;110(9):3585–3590. doi:10.1073/pnas.1216855110.

76. Shriki O, Alstott J, Carver F, Holroyd T, Henson RNA, Smith ML, et al. Neuronal avalanches in the resting MEG of the human brain. Journal of Neuroscience. 2013;33(16):7079–7090. doi:10.1523/jneurosci.4286-12.2013.

77. Massobrio P, Pasquale V, Martinoia S. Self-organized criticality in cortical assemblies occurs in concurrent scale-free and small-world networks. Scientific Reports. 2015;5(1):10578. doi:10.1038/srep10578.

78. Arviv O, Goldstein A, Shriki O. Near-critical dynamics in stimulus-evoked activity of the human brain and its relation to spontaneous resting-state activity. Journal of Neuroscience. 2015;35(41):13927–13942. doi:10.1523/jneurosci.0477-15.2015.

79. Arviv O, Medvedovsky M, Sheintuch L, Goldstein A, Shriki O. Deviations from critical dynamics in interictal epileptiform activity. Journal of Neuroscience. 2016;36(48):12276–12292. doi:10.1523/jneurosci.0809-16.2016.

80. Fekete T, Omer DB, O’Hashi K, Grinvald A, van Leeuwen C, Shriki O. Critical dynamics, anesthesia and information integration: Lessons from multi-scale criticality analysis of voltage imaging data. NeuroImage. 2018;183:919–933. doi:10.1016/j.neuroimage.2018.08.026.

81. Arviv O, Goldstein A, Shriki O. Neuronal avalanches and time-frequency representations in stimulus-evoked activity. Scientific Reports. 2019;9(1):13319. doi:10.1038/s41598-019-49788-5.

82. Priesemann V, Wibral M, Valderrama M, Pröpper R, Le Van Quyen M, Geisel T, et al. Spike avalanches in vivo suggest a driven, slightly subcritical brain state. Frontiers in Systems Neuroscience. 2014;8:108. doi:10.3389/fnsys.2014.00108.

83. Doya K, Ishii S, Pouget A, Rao RPN. Bayesian brain: probabilistic approaches to neural coding. 1st ed. Computational Neuroscience. MIT Press; 2007.

84. Sedley W, Friston KJ, Gander PE, Kumar S, Griffiths TD. An integrative tinnitus model based on sensory precision. Trends in Neurosciences. 2016;39(12):799–812. doi:10.1016/j.tins.2016.10.004.

85. Noda K, Kitahara T, Doi K. Sound change integration error: An explanatory model of tinnitus. Frontiers in Neuroscience. 2018;12:831. doi:10.3389/fnins.2018.00831.

86. Corlett PR, Horga G, Fletcher PC, Alderson-Day B, Schmack K, Powers AR. Hallucinations and strong priors. Trends in Cognitive Sciences. 2019;23(2):114–127. doi:10.1016/j.tics.2018.12.001.

87. Arodz H, Dziarmaga J, Zurek WH. Patterns of symmetry breaking. vol. 127. Springer Science & Business Media; 2003.

88. Dayan P, Abbott LF. Theoretical neuroscience: computational and mathematical modeling of neural systems. Computational Neuroscience. The MIT Press; 2001.

89. Hudspeth A, Corey D. Sensitivity, polarity, and conductance change in the response of vertebrate hair cells to controlled mechanical stimuli. Proceedings of the National Academy of Sciences. 1977;74(6):2407–2411. doi:10.1073/pnas.74.6.2407.

90. Russell I, Sellick P. Intracellular studies of hair cells in the mammalian cochlea. The Journal of Physiology. 1978;284(1):261–290. doi:10.1113/jphysiol.1978.sp012540.

91. Cruickshanks KJ, Wiley TL, Tweed TS, Klein BEK, Klein R, Mares-Perlman JA, et al. Prevalence of hearing loss in older adults in Beaver Dam, Wisconsin: The epidemiology of hearing loss study. American Journal of Epidemiology. 1998;148(9):879–886. doi:10.1093/oxfordjournals.aje.a009713.

92. Agrawal Y, Platz EA, Niparko JK. Prevalence of hearing loss and differences by demographic characteristics among us adults: Data from the national health and nutrition examination survey, 1999-2004. Archives of Internal Medicine. 2008;168(14):1522–1530. doi:10.1001/archinte.168.14.1522.

